# Conspecific larval extracts along with biotoxin to attract-and-kill mosquitoes

**DOI:** 10.1101/596643

**Authors:** Gabriel B. Faierstein, WeiYu Lu, Andréa K. L. S. Sena, Rosângela M. R. Barbosa, Walter S. Leal

**Author notes:** Corresponding author: Walter S. Leal.

## Abstract

One of the strategies of integrated vector management is to lure gravid mosquitoes for surveillance purposes or to entice them to lay eggs in water containing toxins that kill the offspring (attract-and-kill or trap-and-kill). Typically, the major challenge of this approach is the development of a lure that stimulates oviposition plus a toxin with no deterrent effect. *Bacillus thuringiensis* var. *israelensis* (Bti) satisfies the latter criterion, but lures for these autocidal gravid traps are sorely needed. We observed that gravid *Aedes aegypti*, *Ae. albopictus,* and *Culex quinquefasciatus* laid significantly more eggs in cups with extracts from 4th-stage larvae (4L) of the same or different species. No activity was found when 4L were extracted with hexane, diethyl ether, methanol, or butanol, but activity was observed with dimethyl sulfoxide extracts. Larval extracts contained both oviposition stimulant(s)/attractant(s) and deterrent(s), which partitioned in the water and hexane phases, respectively. Lyophilized larval extracts were active after a month, but activity was reduced by keeping the sample at 4°C. In the tested range of 0.1 to 1 larvae-equivalent per milliliter, oviposition activity increased in a dose-dependent manner. In field experiments, *Ae. aegpti* laid significantly more eggs in traps loaded with larval extracts plus Bti than in control traps with water plus Bti.

## Introduction

Integrated vector management is a rational decision-making process to make vector control more efficient, cost effective, ecologically sound, and sustainable. It is ultimately aimed at preventing the transmission of vector-borne diseases^1^. Throughout the world, vector abatement groups (67 agencies in California alone) are constantly engaged in vector surveillance not only to monitor populations of native species and the circulation of pathogens, but also for quarantine of invasive mosquito species (eg, *Aedes (Stegomyia) aegypti* and *Ae. albopictus*) as well as to monitor circulation of new and previously reported pathogens (eg, dengue, chikungunya, Zika, and West Nile viruses). In addition to labor-intensive strategies, such as sampling of immature stages and aspiration of adult mosquitoes from house-to-house, abatement district personnel rely heavily on capturing host- and oviposition-seeking mosquitoes with surveillance traps. Although carbon dioxide is the most effective lure, CO_2_-baited traps capture blood-seeking mosquitoes and thus are less effective for early detection of a pathogen, because they trap many mosquitoes that have never had a blood meal. By contrast, gravid traps are more effective for surveillance, because they target a critical epidemiological stage – the gravid females that imbibed in and digested at least one blood meal and, therefore, are more likely to be infected with a vector-borne pathogen than the general adult population is^2^. Almost all female mosquitoes trapped in gravid traps have had at least one blood meal, which increases the chances of detection of circulating viruses. Additionally, ovitraps can also be used as trap-and-kill systems for direct control of mosquito populations. For a direct trap-and-kill control strategy, ovitraps may be transformed into autocidal gravid ovitraps by adding a biological agent (eg, *Bacillus thuringiensis* var. *israelensis,* Bti), an insecticide, or even an adhesive strip, in addition to a natural or synthetic lure (reviewed in reference^2^).

It has been reported for the last 4 decades that larval-holding water and larval-rearing water are “attractive” to conspecific *Aedes* and *Culex* mosquitoes^3–13^, although it has not been unambiguously determined whether these lures are derived from immature stages of mosquitoes, from bacteria they host, or even from bacteria in the rearing medium. From an evolutionary perspective, the cost-benefit of producing such a signal is intriguing, but from epidemiological and practical viewpoints, it is a weak link worth exploring as a target for vector control. Here, we show that gravid females *Ae. aegypti, Ae. albopictus*, or *Culex quinquefasciatus* mosquitoes lay significantly more eggs in oviposition cups loaded with aqueous extracts from conspecific or allospecific 4th-stage larvae or pupae (but not with extracts from eggs) than in clean water cups. Liquid-liquid extraction of the active larval extracts showed that they contain both oviposition stimulant(s) and deterrent(s) in the aqueous and organic phases, respectively. Field studies in Recife, Brazil showed that *Ae. aegypti* laid significantly more eggs in traps baited with larval extract plus Bti than in traps baited with Bti-containing water, thus demonstrating that the larval extracts are feasible for integrated vector management applications.

## Results & Discussion

Although there is a consensus in the literature that larval- and pupal-holding waters are active in eliciting oviposition in conspecific adult mosquitoes^3–13^, some apparently contradictory results may be derived from confounding factors, such as visual stimuli and overcrowding factors. When evaluating interspecific interactions, the overcrowding factors^14–16^ deserve particular attention. To circumvent these problems, we measured oviposition behavior using 150-ml of water per cup and with standard concentrations of direct extracts from larvae and pupae. Inspired by preliminary and promising experiments with *Ae. albopictus*^10^, we tested extracts at 1 larva-equivalent or 1 pupa-equivalent per 3 ml of water, or 0.33 equivalent per ml. First, we obtained aqueous extracts from L4 larvae of *Ae. aegypti* and tested the fresh extracts against gravid females of *Ae. aegypti*, *Ae. albopictus*, and *Cx. quinquefasciatus.* Both *Ae. aegypti* (Fig. 1A) and *Ae. albopictus* (Fig. 1B) gravid females showed a highly significant preference for cups loaded with *Ae. aegypti* larvae than for control cups (water only), with *Cx. quinquefasciatus* showing a moderate preference (Fig. 1C). Likewise, gravid females of the 3 species laid significantly more eggs in cups containing aqueous extract from *Ae. aegypti* pupae than in water cups (Fig. 1D-E). By contrast, none of the 3 species showed oviposition preference for aqueous extracts from *Ae. aegypti* eggs (Fig. S1). Our findings differ from what has been reported, ie, that responses of gravid *Ae. aegypti* females to conspecific larval rearing water did not differ significantly from water controls^3,12^. This discrepancy may be due to the difference in extracts (larval rearing water vs direct extract) or loss of activity of larval rearing water over a short period of time. We tested the longevity of our extracts later, but first asked whether the oviposition attractant/stimulant could also be extracted from *Ae. albopictus.* Again, gravid females of the tested species laid significantly more eggs in cups loaded with *Ae. albopictus* larval extracts than in control cups (Figs. 2A-C). Interestingly, the pupal extracts from *Ae. albipictus* were active against conspecific and *Cx. quinquefasciatus* adult females (Fig. 2D, F), but not against *Ae. aegypti* (Fig. 2E). *Ae. albopictus* showed a preference for conspecific egg extracts over control water cups (Fig. S2), but *Ae. aegypti* and *Cx. quinquefasciatus* did not. These findings are somewhat consistent with earlier preliminary experiments showing that extracts from *Ae. albopictus* larvae and pupae (but not eggs) were active to conspecific gravid females^10^. We next tested extracts from *Cx. quinquefasciatus* L4 larvae and pupae. Again, gravid females of the 3 mosquito species laid significantly more eggs in cups loaded with larval extract than in control cups (Fig. 3A-C) as well as in cups loaded with extracts from *Cx. quinquefasciatus* pupae than in plain water cups (Fig. 3D-F). Although it is tempting to assume that larval extracts from *Ae. aegypti, Ae. albopictus*, and *Cx. quinquefasciatus* share common active ingredient(s), this assumption must await further rigorous testing and identification of the active ingredient(s) of these extracts.

**Figure 1.**
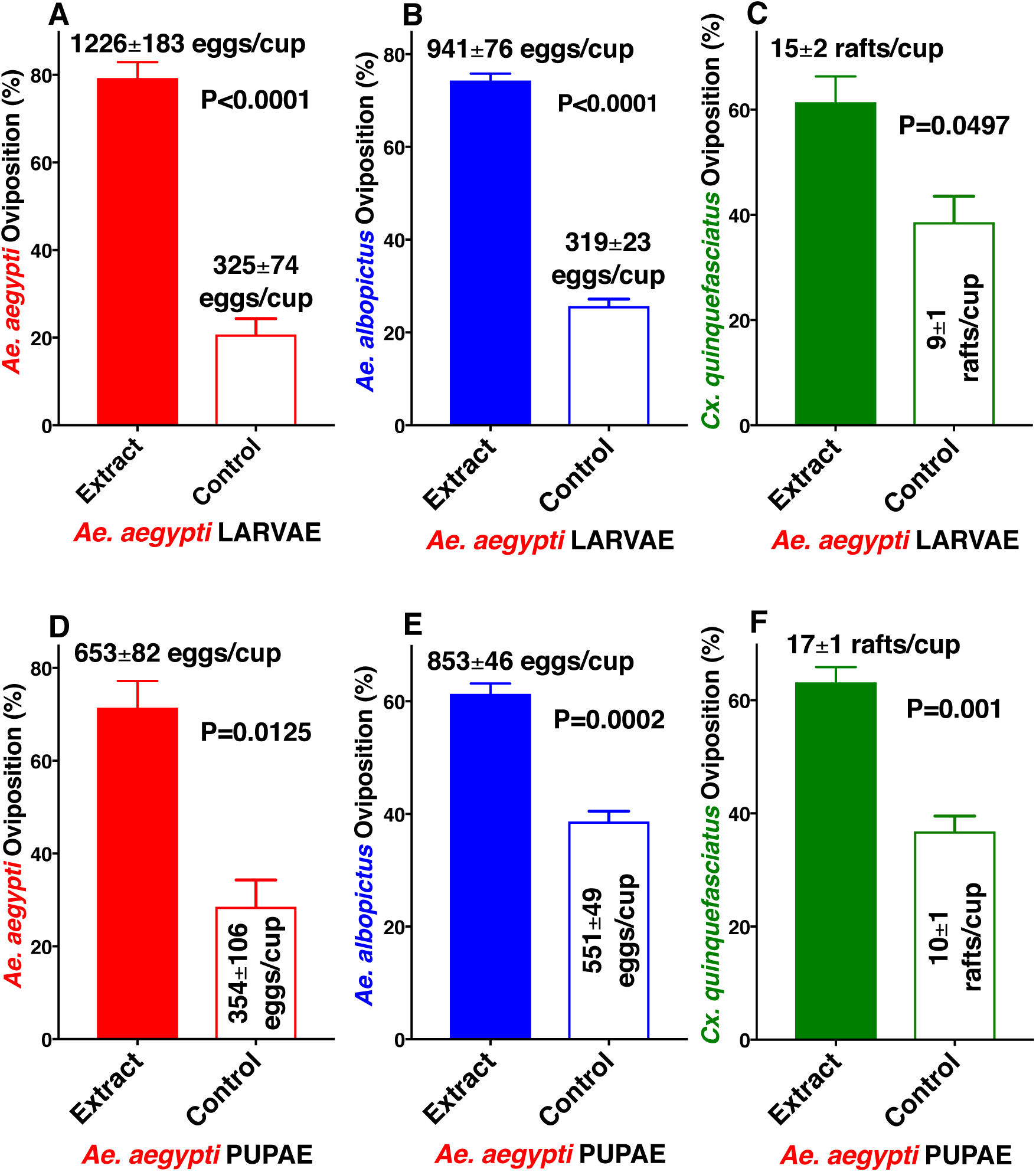
Oviposition preference by *Ae. aegypti*, *Ae. albopictus,* and *Cx. quinquefasciatus* to aqueous extracts from *Ae. aegypti* larvae or pupae compared with water. Mean (±SEM) number of eggs laid by (A) *Ae. aegypti* and (B) *Ae. albopictus*, and egg rafts laid by (C) *Cx. quinquefasciatus* in cups loaded with *Ae. aegypti* 4th-stage larval extracts and control cups (water only). (D, E, F) Oviposition preference by the same species when given a choice of *Ae. aegypti* pupal extracts and water only. N = 10 for each treatment. For clarity, data are presented in percentage of oviposition preference, with mean number of eggs or egg rafts presented along with each bar. After arcsine transformation and passing the Shapiro-Wilk normality test, each dataset was compared using the 2-tailed, paired *t* test.

**Figure 2.**
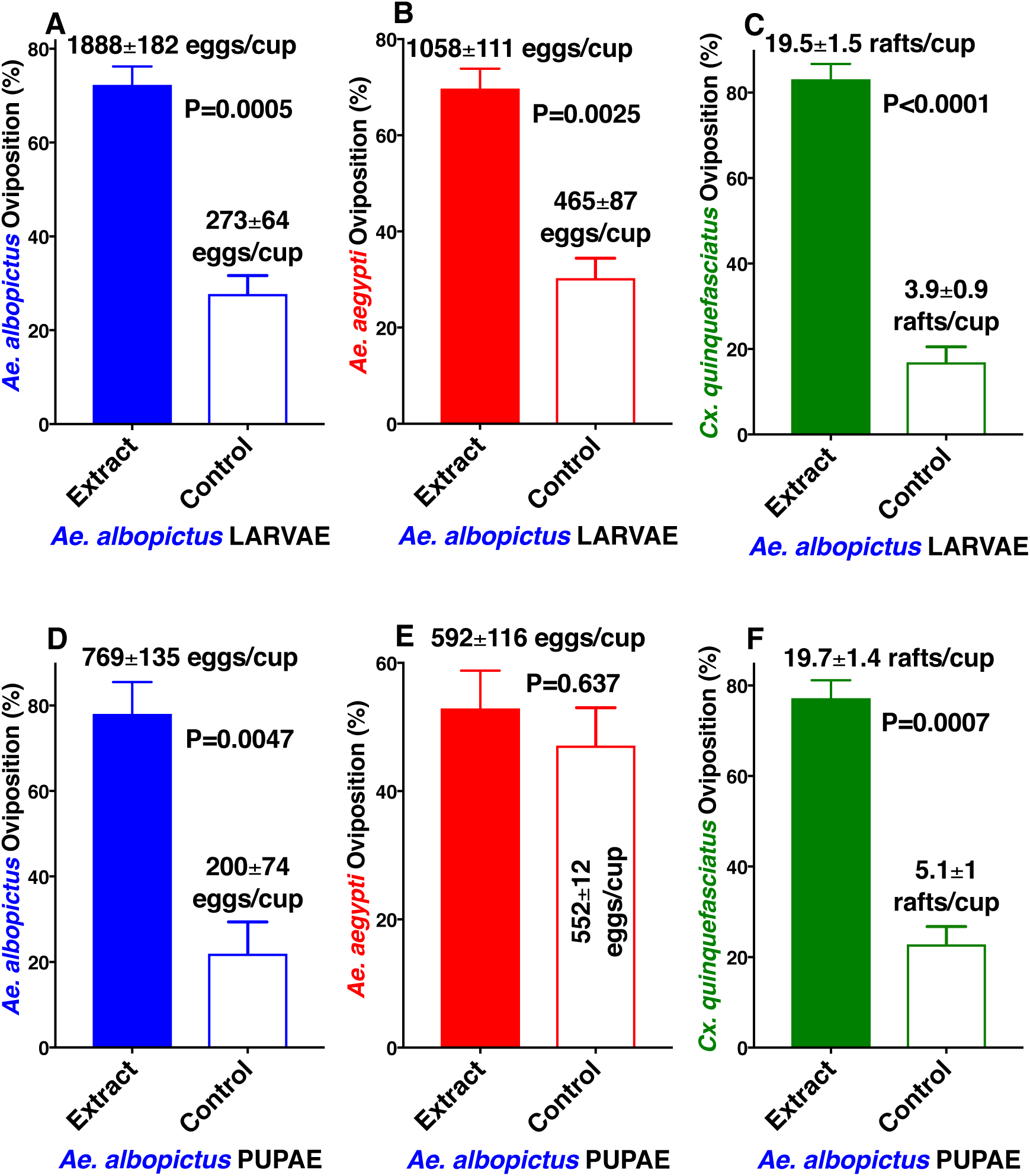
Oviposition preference by *Ae. albopictus, Ae. aegypti*, and *Cx. quinquefasciatus* to aqueous extracts from *Ae. albopictus* larvae or pupae compared with water. Mean (±SEM) number of eggs laid by (A) *Ae. albopictus* and (B) *Ae. aegypti*, and egg rafts laid by (C) *Cx. quinquefasciatus* in cups loaded with *Ae. albopictus* 4th-stage larval extracts and control cups (water only). (D, E, F) Oviposition preference by the same 3 species in dual choices assays comparing *Ae. albopictus* pupal extracts and water only. N = 10 for each treatment. For clarity, data are presented in percentage of oviposition preference, with mean number of eggs or egg rafts presented along with each bar. After arcsine transformation and passing the Shapiro-Wilk normality test, each dataset was compared by using the 2-tailed, paired *t* test.

**Figure 3.**
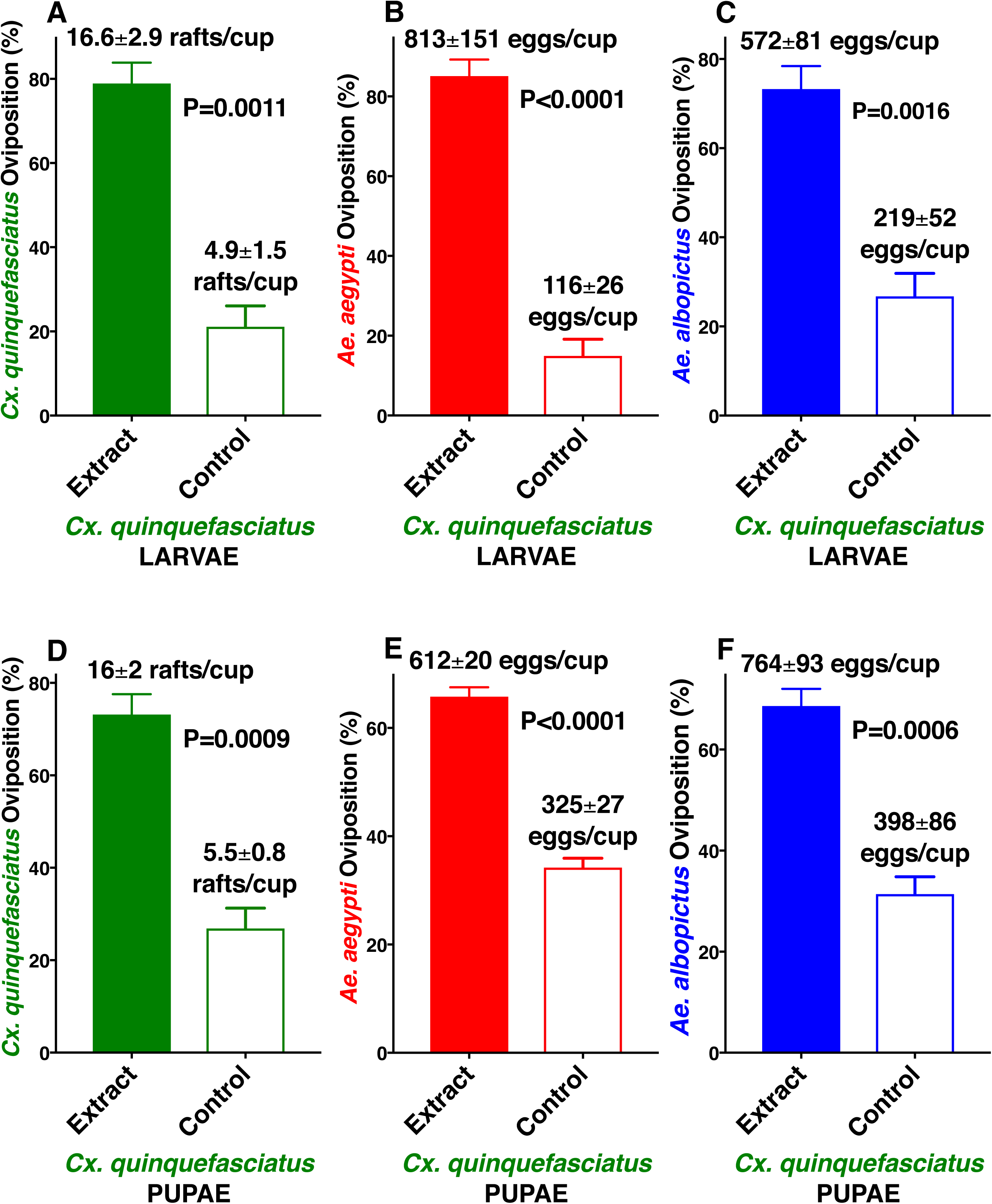
Oviposition preference by *Cx. quinquefasciatus, Ae. aegypti*, and *Ae. albopictus* to aqueous extracts from *Cx. quinquefasciatus* larvae or pupae compared with water. Mean (±SEM) number of egg rafts laid by (A) *Cx. quinquefasciatus*, and eggs laid by (B) *Ae. aegypti* and (C) *Ae. albopictus* in cups loaded with *Cx. quinquefasciatus* 4th-stage larval extracts and control cups (water only). (D, E, F) Oviposition preference by the same 3 species in dual choices assays comparing *Cx. quinquefasciatus* pupal extracts and water only. N = 10 for each treatment. For clarity, data are presented in percentage of oviposition preference, with the mean number of eggs or egg rafts presented along with each bar. After arcsine transformation and passing the Shapiro-Wilk normality test, each dataset was compared by using the 2-tailed, paired *t* test.

A very hydrophobic compound, n-heneicosane^17^, has been isolated from *Ae. aegypti* eggs and has been demonstrated to stimulate the antennae^18^ of both *Ae. aegypti* and *Ae. albopictus* and thus has been suggested to be an oviposition attractant^17,18^. Although it is highly unlikely that n-heneicosane would be an active ingredient in the larval and pupal extracts, we tested whether the active ingredients could be extracted with organic solvents. To avoid emulsification when the extracts were mixed with water in oviposition cups, hexane extracts were dried up and reconstituted in dimethyl sulfoxide (DMSO). Indeed, there was no significant difference in the number of eggs laid by *Ae. aegypti* gravid females in cups loaded with hexane extract vs control cups (Fig. 4A). By contrast, there was a significant preference for cups loaded with DMSO larval extracts compared with the control (water plus DMSO) (Fig. 4B). Similarly, *Cx. quinquefasciatus* showed a significant preference for DMSO but not for hexane extracts (Fig. 4C, D). We repeated these experiments and noticed a trend of controls getting more egg rafts than hexane extracts, thus suggesting a possible deterrent effect from hexane extracts. We surmised that a trace of these or other deterrents might be contained in our aqueous extracts. To test this assumption, we performed liquid-liquid extraction of the active material and tested separately the aqueous and organic phases. Of note, a small gel-like intermediate phase was discarded after the aqueous phase was collected and before the start of collecting the hexane phase. There was a clear preference for gravid *Cx. quinquefasciatus* to lay eggs in the aqueous fraction over the control (Fig. 5A), whereas the organic phase showed a deterrent effect (Fig. 5B). We, therefore, concluded that the aqueous extracts contain both oviposition stimulant(s)/attractant(s) and deterrent(s) with the former offsetting the latter. We then extracted *Cx. quinquefasciatus* L4 larvae with other organic solvents and found similar deterrent effects with diethyl ether, methanol, and butanol (Fig. S3). Furthermore, we surmised that the active ingredient is either water soluble organic compound(s) or protein(s)/peptide(s) that do not require folding for activity; otherwise activity in DMSO extracts would have been lost^19^.

**Figure 4.**
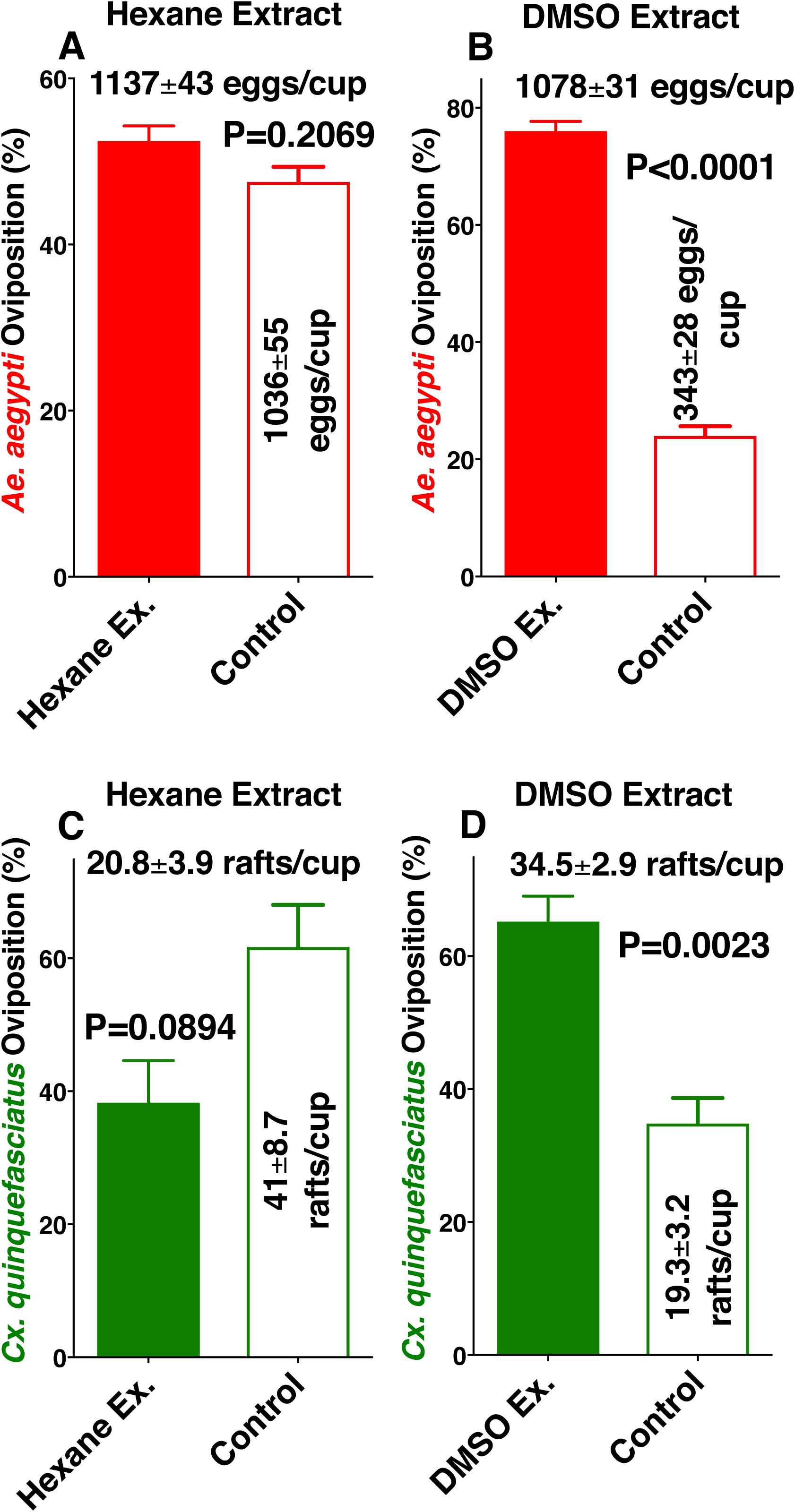
Oviposition preference by the yellow fever mosquito and the southern house mosquito to conspecific larval extracts with hexane or DMSO. Oviposition preference by *Ae. aegypti* comparing (A) hexane and (B) DMSO extracts from conspecific 4th-stage larvae vs water. Mean (±SEM) number of egg rafts laid by *Cx. quinquefasciatus* in cups loaded with conspecific 4th-stage larval extracts obtained with (C) hexane or (D) DMSO and water cups.

**Figure 5.**
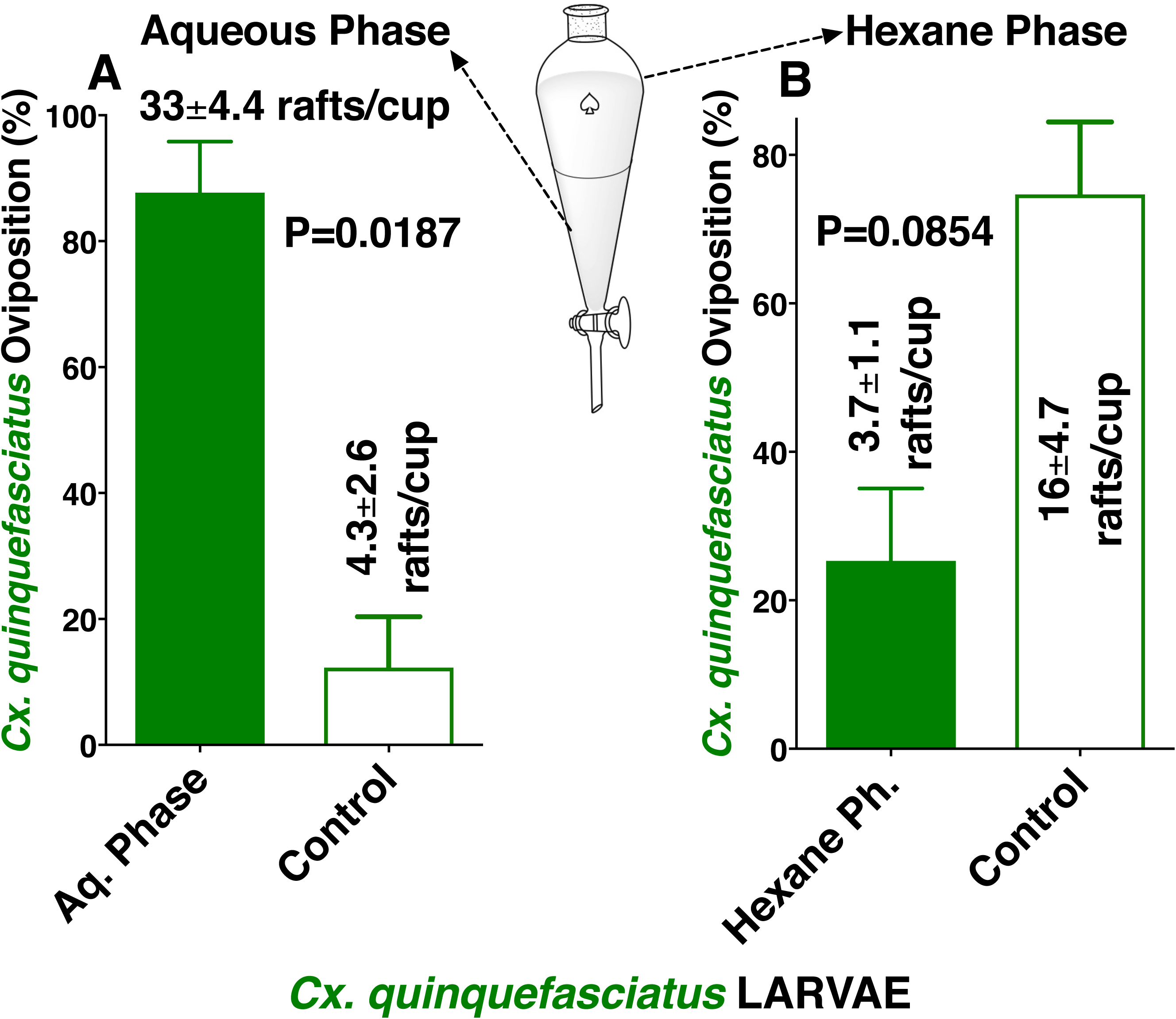
Evidence for oviposition stimulant(s) and deterrent(s) in extracts from 4th-stage *Cx. quinquefasciatus*. Aqueous larval extract was partitioned with hexane, and subsequently the 2 phases were tested for oviposition preference, ie, (A) aqueous phase and (B) hexane phase. To avoid emulsion formation, hexane extract was dried, and the solvent replaced with DMSO. An equal amount of DMSO was added to the control cup. N = 4 for each treatment. After arcsine transformation and passing the Shapiro-Wilk normality test, each dataset was compared by using the 2-tailed, paired *t* test.

Next, we investigated whether lyophilization would affect activity. Larval extracts from the yellow fever mosquito were separated into 2 groups; half of the sample was extracted and then kept at 4°C for 3 days, and the other half of the sample was lyophilized and 3 days later extracted just before bioassays. Responses elicited by the refrigerated and lyophilized samples were significantly higher than in their respective controls (Fig. 6A, B). Interestingly, however, when these experiments were performed with a longer storage time (30 days), the refrigerated sample lost activity, whereas activity was retained by the lyophilized sample (Fig. 6C, D). These experiments reinforce what has been observed with direct organic solvent extractions. Specifically, it is highly unlikely that the active ingredients are organic molecules of low or medium molecular weight, which would have evaporated during lyophilization. Moreover, these data show that the active ingredient(s) undergoes degradation at 4°C as would be expected for a peptide or protein kept in a crude extract, which must contain proteolytic enzymes from the mosquito gut.

**Figure 6.**
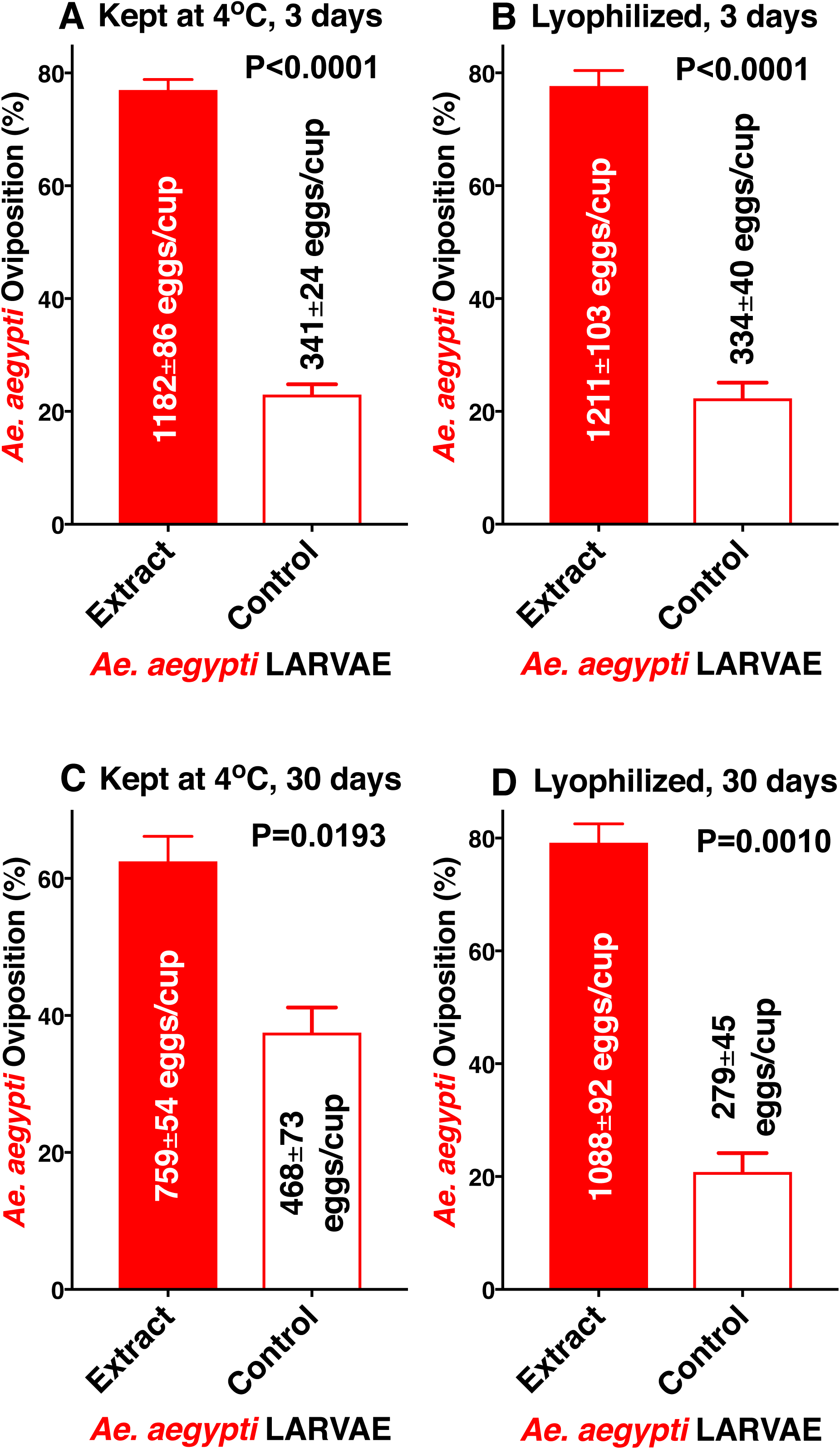
Assessing stability of the larvae-derived oviposition stimulant(s). *Ae. aegypti* oviposition preference to conspecific larval extracts (A) kept at 4°C for 3 days and (B) freshly lyophilized, kept at room temperature and reconstituted 3 days later. Similar experiments performed with fresh extract (C) kept at 4°C for 30 days and (D) freshly lyophilized extract kept at room temperature for 30 days and reconstituted on the day of the tests. N = 12 for each dataset. Data were arcsine transformed and after passing the Shapiro-Wilk normality test, each group was compared by using 2-tailed, paired *t* tests.

It is very common in chemical ecology that some compounds act in a dose-dependent manner, being an attractant at lower doses and a deterrent at higher doses. Because we used a standard concentration of 0.33 L-eq/ml throughout these studies, we next tested lower and higher doses. The activity from 0.1-1 L-eq/ml increased in a dose-dependent manner (Fig. 7). It is therefore unlikely that the oviposition stimulant(s)/attractant(s) in our aqueous extracts are related to overcrowding factors. The active lures are like exudates from larvae (and pupae), but we cannot unambiguously determine whether they are derived from bacteria housed in mosquito gut or by the insect. From an evolutionary perspective, it would make more sense if they were produced by bacteria rather than by the host.

**Figure 7.**
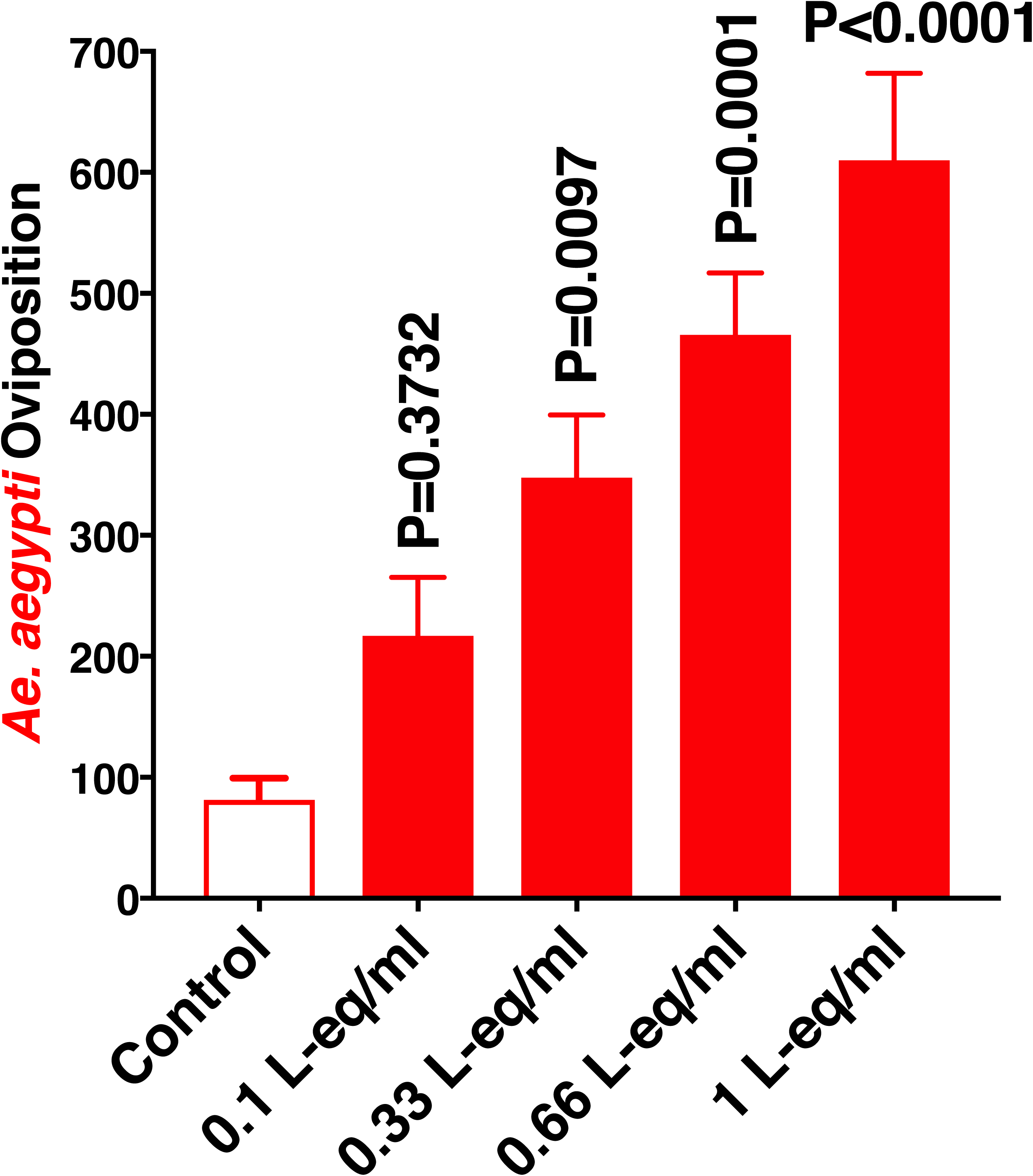
Effect of the concentration of larval extracts on oviposition stimulation. Larval extracts from 4th-stage *Ae. aegypti* were tested in indoor assays comparing the “standard dilution” of 0.33 larvae-equivalent per ml (L-eq/ml) with lower and higher doses. N = 12. Means of the treatments were compared with the control by using the nonparametric Friedman test.

Lastly, we explored the potential application of these larval extracts in attraction-and-kill strategies. Specifically, we questioned whether these extracts would be active in the field when combined with a toxic agent. The number of eggs in traps loaded with both larval extract and Bti were significantly higher than in the control traps with water plus Bti (Fig. 8). In conclusion, L4 larval extracts have a potential application in integrated vector management. The logistics of this attract-and-kill strategy might be simplified when the active ingredients are identified and synthetic counterparts are used instead of cumbersome crude extracts. For the time being, however, extracts from lyophilized larvae may be used as lure.

**Figure 8.**
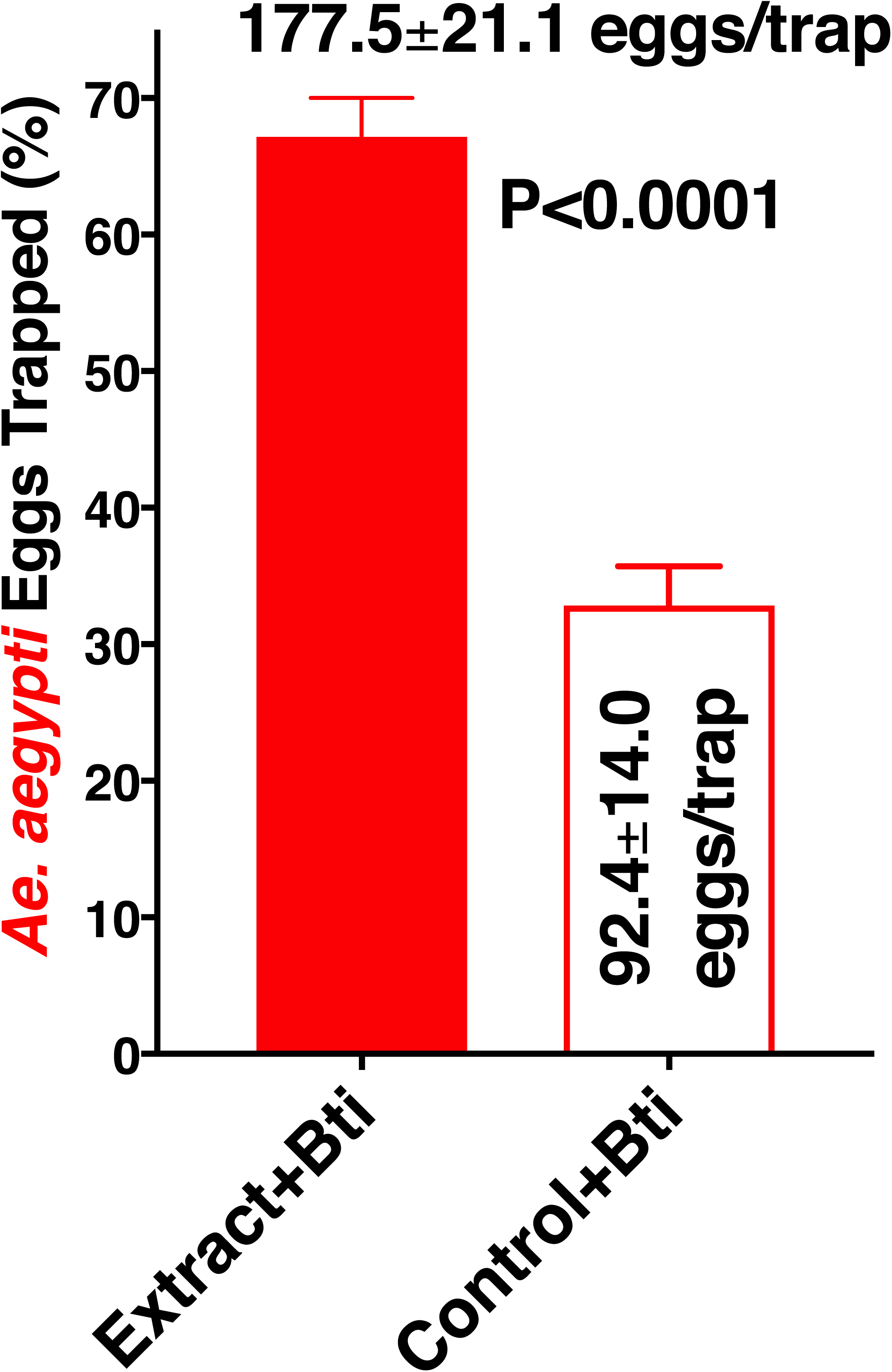
Oviposition preference for larval extracts in the presence of *Bacillus thuringiensis israelensis* (Bti). Bti was added to traps loaded with 4th-stage larval extracts from *Ae. aegypti* as well as to the control water traps. Pairs of traps were deployed in the 8 different locations in the field and inspected every 2 weeks. N = 51. Means were compared by using the Wilcoxon matched-pairs signed rank test.

## Materials & Methods

### Mosquitoes

*Cx. quinquefasciatus* mosquitoes used in this study at UC Davis originated from Dr. Anthon Cornel’s stock laboratory colony, which in turn started from adult mosquitoes collected in Merced, CA, in the 1950s. The Davis colony has been kept for more than 7 years at 27±1°C, 75%±5 relative humidity, and under a photoperiod of 12:12 h (light:dark). The Recife colony of *Cx. quinquefasciatus* originated from eggs collected in Peixinhos, a neighborhood of Olinda, metropolitan region of Recife, Pernambuco, Brazil in 2009. The *Ae. aegypti* and *Ae. albopictus* colonies started in 1996 and 1998, respectively, from eggs collected in neighborhoods in Recife. All 3 mosquito colonies from Brazil were kept in Recife at 26±2°C, 65-85% relative humidity, and under a photoperiod of 12:12 h (light:dark). Larvae were kept in plastic containers (30 × 15 cm; 10 cm height) with a density of approximately 0.3 larvae/ml.

### Extraction procedures

Fourth-stage larvae were collected with a plastic mesh net and washed with distilled water 3-7 times. Fifty larvae were placed into a 2-ml microcentrifuge tube. After adding 0.5 ml of distilled water, the larvae were grinded, the pistil was washed twice with 0.5 ml of distilled water. The extract was then filtered through a Whatman #1 filter paper (catalogue number 1001-110) and washed with a total 150 ml of distilled water. Organic solvent extracts followed a slightly different procedure. Hexane, diethyl ether, methanol, and butanol extracts were obtained in Pyrex glass homogenizers, the supernatant was filtered through Pasteur pipettes with a cotton plug, and this procedure was repeated twice. In the case of hexane and diethyl ether extracts, after separation the solvent was evaporated, and the extract reconstituted in DMSO. Methanol and butanol extracts were used directly without solvent exchange. DMSO extracts were centrifuged to remove debris. For experiments comparing lyophilization with refrigeration, a group of L4 larvae was separated into 2 equal parts; 1 sample was directly extracted with water and the other was lyophilized before extraction. For partition with hexane, a freshly prepared aqueous extract was transferred to a separatory funnel and equal volume of hexane was added. After vigorous shaking, the 2 phases were separated. The concentration of the aqueous phase was adjusted, a small gel-like intermediate phase was discarded, the organic phase was dried up in a rotavapor, reconstituted in DMSO, and the concentration was adjusted.

### Oviposition bioassay

This bioassay was performed in cages (50 x 40 x 32 cm) in which 2 oviposition cups were placed in diagonal positions 30 cm away from each other^20^. These cups were loaded one with treatment and another with control. In both cases, the volume was adjusted to 150 ml with water. The 2 cups had the same amount of solvent. For example, the same amounts of methanol, butanol, and DMSO were added to both cups to deliver the larvae-equivalent to make a final dose of 0.33 L-eq/ml and to have the same amount of solvent in control cups. Twelve cages were used per treatment per day. For experiments with *Aedes* mosquitoes, we used a filter paper on the edge of the cups as an oviposition substrate. This was not necessary with *Cx. quinquefasciatus* because they lay eggs on the water edge. Thirty to 50 gravid females were released per cage. Egg rafts from *Cx. quinquefasciatus* were collected daily, whereas eggs from *Aedes* mosquitoes were counted after 7 days. Data were analyzed with Prism 7 (GraphPad, La Jola, CA). They were arcsine transformed, and after passing the Shapiro-Wilk normality test, they were compared by using the 2-tailed, paired *t* test. For comparison of doses, 4 treatments and 1 control were placed in each case. Thus, means of treatments were compared with the control by using the nonparametric Friedman test.

### Field experiments

These experiments were performed on the campus of the Federal University of Pernambuco. Ovitraps^21^ were loaded with 1 liter of larval extract (final dose, 0.33 L-eq/ml) plus 0.5 g of *Bacillus thuringiensis* serotype *israelensis* (VectorBac® WG, strain AM65-62, Lot: 257-352-PG), whereas the control traps were loaded with 1 liter of water and 0.5 g of Bti. To each trap, 2 wood boards (5 x 15 cm; 5 mm thickness) were attached to the border of the water container so as to facilitate oviposition. These experiments were performed from October 2017 to February 2018. Traps were inspected and rotated every 2 weeks, with a total of 51 replicates. Data were analyzed by comparing the means by using the Wilcoxon matched-pairs signed rank test.

## Supporting information

Raw data (dataset 1)

## Data Availability

All raw data are included, see Supplemental Information.

## Acknowledgements

We thank Dr. Anthon Cornel (UC Davis) for providing mosquitoes that were used to duplicate his *Cx. quinquefasciatus* colony at the Davis campus. This work was supported by grants from the National Institutes of Health R01AI095514 and R21AI128931.

## Author Contributions

W.S.L. and R.M.R.B. designed the research; G.B.F., W.L., A.K.L.S.S. and W.S.L performed the extraction; G.B.F., W.L. and A.K.L.S.S. performed behavioral assays; G.B.F., W.L., R.M.R.B. and W.S.L. analyzed the data; W.S.L. prepared the figures and wrote the manuscript. All authors critically reviewed and approved the final version of the manuscript.

## Additional Information

**Supplementary information** accompanies this paper at .....

## Supplementary Information

**Figure S1.**
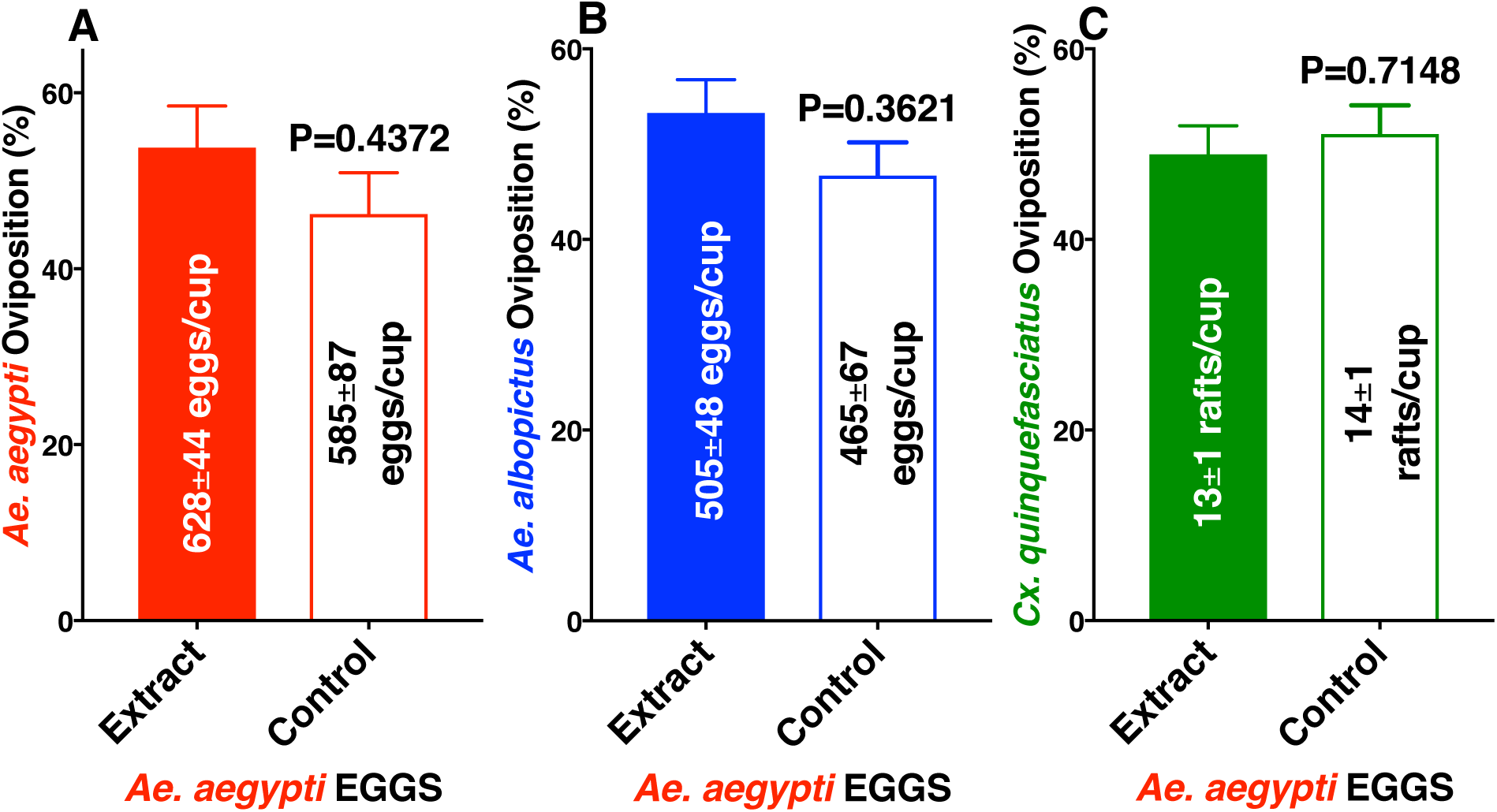
Oviposition preference by *Ae. aegypti*, *Ae. albopictus,* and *Cx. quinquefasciatus* to aqueous extracts from eggs of *Ae. aegypti* compared with water. Mean (±SEM) number of eggs laid by (A) *Ae. aegypti* and (B) *Ae. albopictus*, and egg rafts laid by (C) *Cx. quinquefasciatus* in cups loaded with extracts and control cups (water only). N = 10 for each treatment. For clarity, data are presented in percentage of oviposition preference, with mean number of eggs or egg rafts presented along with each bar. After arcsine transformation and passing the Shapiro-Wilk normality test, each dataset was compared by using the 2-tailed, paired *t* test.

**Figure S2.**
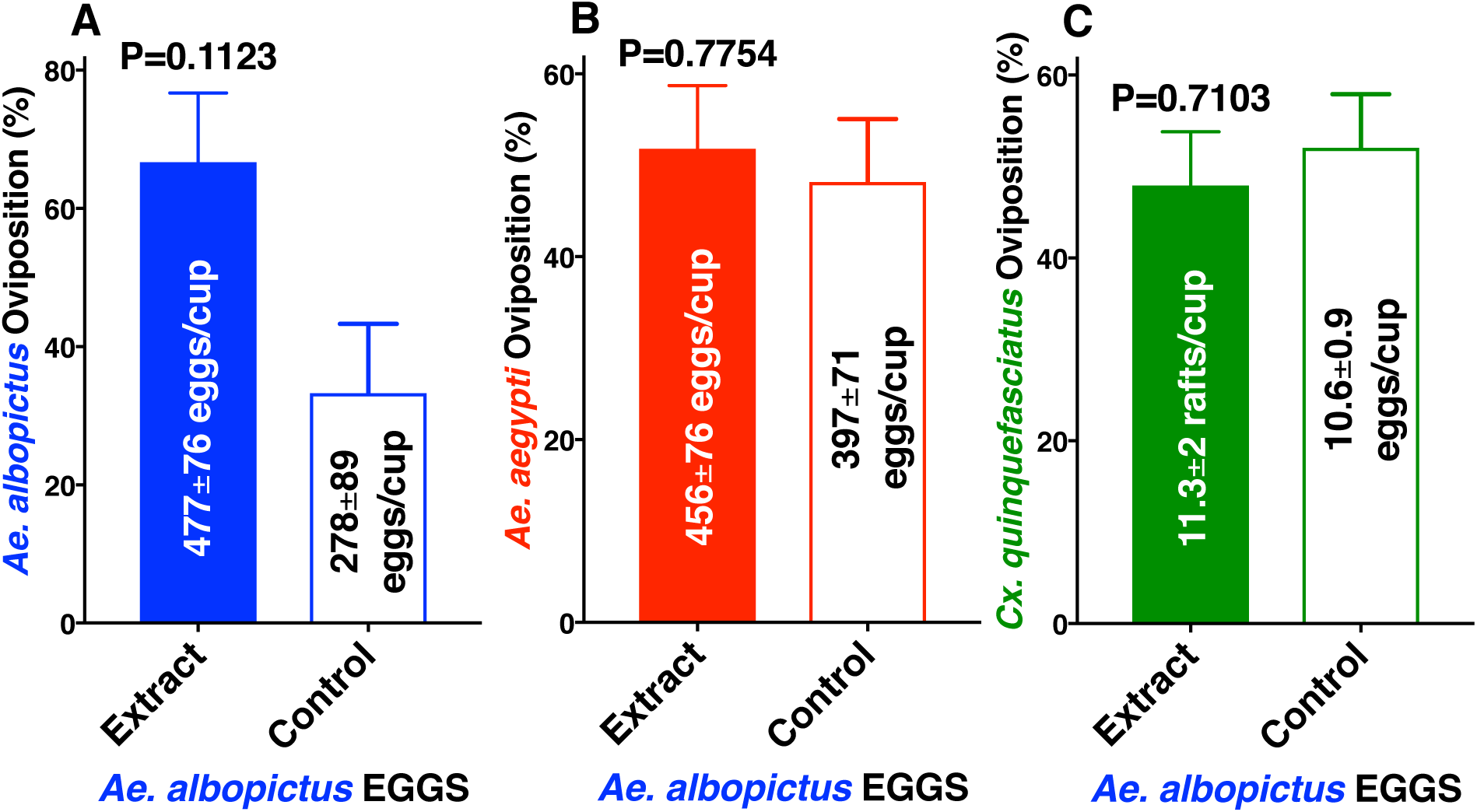
Oviposition preference by *Ae. albopictus, Ae. aegypti*, and *Cx. quinquefasciatus* to aqueous extracts from *Ae. albopictus* eggs compared with water. Mean (±SEM) number of eggs laid by (A) *Ae. albopictus* and (B) *Ae. aegypti*, and egg rafts laid by (C) *Cx. quinquefasciatus* in cups loaded with *Ae. albopictus* egg extracts and control cups (water only). (D, E, F) Oviposition preference by the same 3 species in dual choices assays comparing *Ae. albopictus* pupal extracts and water only. N = 10 for each treatment. For clarity, data are presented in percentage of oviposition preference, with mean number of eggs or egg rafts presented along with each bar. After arcsine transformation and passing the Shapiro-Wilk normality test, each dataset was compared by using the 2-tailed, paired *t* test.

**Figure S3.**
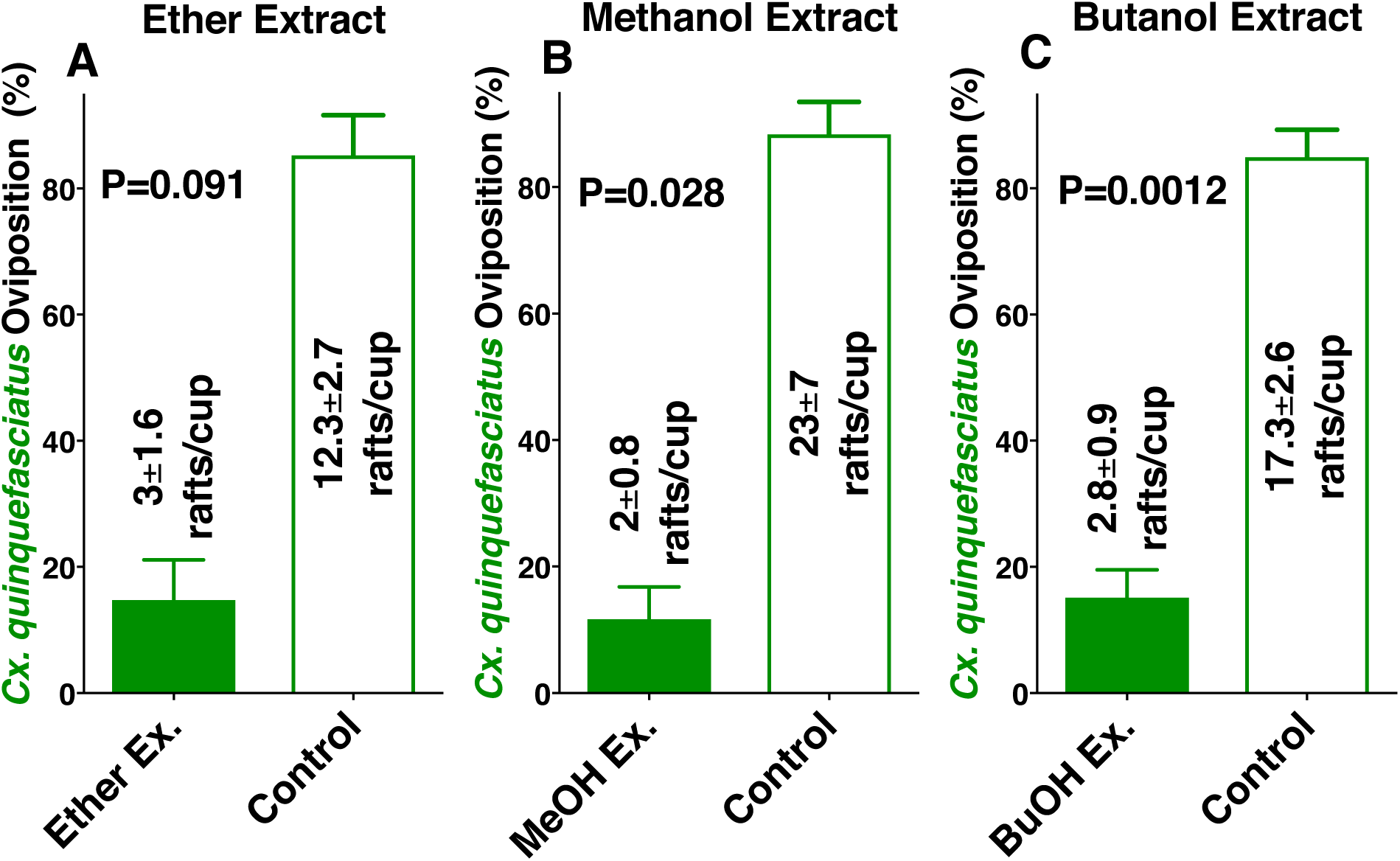
Oviposition preference by *Cx. quinquefasciatus* in dual choice assays comparing water vs conspecific larval extracts with various solvents. Percentage of oviposition preference comparing control traps with those loaded with (A) diethyl ether (ether), (B) methanol, and (C) butanol extracts from 4th-stage *Cx. quinquefasciatus* larvae. N = 12 for each treatment. After arcsine transformation and passing the Shapiro-Wilk normality test, each dataset was compared by using the 2-tailed, paired *t* test.

