## Supplementary material for "Conspecific larval extracts along with biotoxin to attract-and-kill mosquitoes": Raw data (dataset 1)

**Raw Data for Figure 1**

| Ae. aegypti | | Ae. albopictus | | Cx. quinquefasciatus | |
|---|---|---|---|---|---|
| Extract | Control | Extract | Control | Extract | Control |
| 419 | 32 | 791 | 223 | 18 | 9 |
| 1899 | 246 | 1266 | 306 | 8 | 14 |
| 1561 | 583 | 879 | 241 | 24 | 4 |
| 1549 | 54 | 1203 | 320 | 24 | 4 |
| 630 | 163 | 607 | 251 | 16 | 10 |
| 790 | 440 | 863 | 291 | 5 | 4 |
| 1417 | 309 | 721 | 329 | 13 | 9 |
| 1572 | 343 | 837 | 391 | 10 | 12 |
| 1916 | 793 | 921 | 408 | 18 | 10 |
| 504 | 291 | 1319 | 429 | 15 | 13 |

**Raw Data for Figure 2**

| Ae. albopictus | | Ae. aegypti | | Cx. quinquefasciatus | |
|---|---|---|---|---|---|
| Extract | Control | Extract | Control | Extract | Control |
| 878 | 551 | 850 | 558 | 22 | 3 |
| 1888 | 616 | 1287 | 92 | 10 | 4 |
| 498 | 155 | 807 | 563 | 23 | 2 |
| 231 | 183 | 1463 | 636 | 25 | 1 |
| 602 | 416 | 1357 | 1001 | 21 | 4 |
| 106 | 70 | 1623 | 133 | 19 | 4 |
| 475 | 45 | 843 | 369 | 17 | 4 |
| 509 | 94 | 518 | 262 | 25 | 3 |
| 1626 | 307 | 878 | 383 | 17 | 2 |
| 869 | 293 | 951 | 653 | 16 | 12 |

**Raw Data for Figure 3**

| Cx. quinquefasciatus | | Ae. aegypti | | Ae. albopictus | |
|---|---|---|---|---|---|
| Extract | Control | Extract | Control | Extract | Control |
| 2 | 1 | 302 | 90 | 331 | 274 |
| 15 | 1 | 814 | 20 | 796 | 147 |
| 28 | 2 | 1171 | 173 | 298 | 321 |
| 28 | 2 | 1524 | 230 | 199 | 40 |
| 2 | 0 | 1589 | 54 | 536 | 80 |
| 18 | 4 | 274 | 253 | 589 | 119 |
| 13 | 8 | 484 | 84 | 874 | 331 |
| 17 | 14 | 419 | 32 | 558 | 574 |
| 22 | 8 | 819 | 160 | 996 | 246 |
| 21 | 9 | 730 | 68 | 544 | 62 |

**Raw Data for Figure 4**

| Ae. aegypti | | | | Cx. quinquefasciatus | | | |
|---|---|---|---|---|---|---|---|
| Hexane Ex. | Control | DMSO Ex. | Control | Hexane Ex. | Control | DMSO Ex. | Control |
| 1346 | 662 | 1221 | 421 | 18 | 25 | 24 | 34 |
| 929 | 847 | 1165 | 440 | 12 | 25 | 31 | 31 |
| 1329 | 1021 | 1067 | 235 | 14 | 15 | 34 | 8 |
| 1116 | 1094 | 998 | 430 | 15 | 46 | 37 | 14 |
| 1274 | 981 | 1093 | 294 | 34 | 16 | 38 | 22 |
| 874 | 1217 | 985 | 426 | 51 | 22 | 35 | 8 |
| 1120 | 1096 | 1139 | 262 | 3 | 59 | 49 | 20 |
| 1121 | 1094 | 1093 | 298 | 3 | 105 | 41 | 42 |
| 1206 | 983 | 1194 | 206 | 31 | 83 | 44 | 16 |
| 1047 | 871 | 846 | 397 | 29 | 63 | 44 | 14 |
| 1228 | 1374 | 981 | 235 | 25 | 18 | 21 | 10 |
| 1051 | 1197 | 1149 | 474 | 15 | 15 | 16 | 13 |

**Raw Data for Figure 5**

| Aq. Phase | Control | Aq. Phase | Control |
|---|---|---|---|
| 32 | 1 | 32 | 1 |
| 38 | 3 | 38 | 3 |
| 41 | 1 | 41 | 1 |
| 21 | 12 | 21 | 12 |

**Raw Data for Figure 6**

| 3 days | | | | 30 days | | | |
|---|---|---|---|---|---|---|---|
| 4oC | | Lyophilized | | 4oC | | Lyophilized | |
| Extract | Control | Extract | Control | Extract | Control | Extract | Control |
| 1286 | 299 | 1223 | 479 | 840 | 336 | 1094 | 289 |
| 1234 | 451 | 1662 | 156 | 657 | 362 | 893 | 380 |
| 1336 | 276 | 1266 | 532 | 872 | 793 | 953 | 314 |
| 1334 | 201 | 1240 | 174 | 553 | 542 | 1487 | 94 |
| 1454 | 363 | 1738 | 173 | 884 | 452 | 912 | 214 |
| 1502 | 414 | 1683 | 259 | 749 | 321 | 1188 | 384 |
| 1237 | 431 | 839 | 477 | | | | |
| 550 | 327 | 906 | 357 | | | | |
| 1384 | 373 | 1203 | 309 | | | | |
| 969 | 424 | 1297 | 485 | | | | |
| 684 | 207 | 817 | 406 | | | | |
| 1208 | 329 | 659 | 198 | | | | |

**Raw Data for Figure 7**

| Control | 0.1 L-eq/ml | 0.33 L-eq/ml | 0.66 L-eq/ml | 1 L-eq/ml |
|---|---|---|---|---|
| 26 | 187 | 414 | 571 | 584 |
| 12 | 78 | 485 | 679 | 711 |
| 79 | 188 | 122 | 556 | 270 |
| 105 | 105 | 357 | 492 | 928 |
| 174 | 132 | 280 | 511 | 930 |
| 19 | 42 | 543 | 89 | 847 |
| 100 | 282 | 340 | 660 | 208 |
| 73 | 128 | 131 | 178 | 547 |
| 192 | 433 | 131 | 493 | 712 |
| 0 | 357 | 699 | 406 | 480 |
| 129 | 591 | 235 | 551 | 772 |
| 67 | 81 | 434 | 404 | 332 |

**Raw Data for Figure 8**

| Control+Bti | Extract+Bti | | Control+Bti | Extract+Bti |
|---|---|---|---|---|
| 22 | 1 | | 80 | 110 |
| 73 | 2 | | 29 | 59 |
| 72 | 176 | | 40 | 146 |
| 1 | 90 | | 125 | 206 |
| 37 | 129 | | 0 | 57 |
| 43 | 58 | | 57 | 71 |
| 127 | 291 | | 0 | 127 |
| 585 | 443 | | 156 | 193 |
| 28 | 51 | | 48 | 103 |
| 45 | 59 | | 153 | 232 |
| 0 | 120 | | 42 | 136 |
| 130 | 157 | | 197 | 95 |
| 0 | 47 | | 102 | 122 |
| 44 | 231 | | 9 | 47 |
| 256 | 682 | | 92 | 203 |
| 167 | 610 | | 75 | 173 |
| 36 | 60 | | 217 | 262 |
| 3 | 4 | | 33 | 50 |
| 274 | 466 | | 130 | 299 |
| 115 | 237 | | 217 | 416 |
| 46 | 161 | | 192 | 441 |
| 130 | 157 | | 31 | 111 |
| 0 | 47 | | 150 | 389 |
| 44 | 231 | | | |
| 108 | 116 | | | |
| 97 | 115 | | | |
| 48 | 65 | | | |
| 5 | 201 | | | |

**Raw Data for Figure S1**

| Ae. aegypti | | Ae. albopictus | | Cx. quinquefasciatus | |
|---|---|---|---|---|---|
| Extract | Control | Extract | Control | Extract | Control |
| 637 | 484 | 234 | 181 | 9 | 12 |
| 723 | 634 | 561 | 484 | 13 | 15 |
| 568 | 963 | 527 | 634 | 17 | 12 |
| 831 | 245 | 596 | 663 | 12 | 10 |
| 354 | 857 | 697 | 245 | 8 | 22 |
| 697 | 755 | 587 | 669 | 15 | 14 |
| 586 | 478 | 496 | 755 | 13 | 16 |
| 468 | 214 | 633 | 478 | 14 | 12 |
| 713 | 896 | 466 | 214 | 18 | 13 |
| 698 | 325 | 258 | 325 | 15 | 14 |

**Raw Data for Figure S2**

| Ae. albopictus | | Ae. aegypti | | Cx. quinquefasciatus | |
|---|---|---|---|---|---|
| Extract | Control | Extract | Control | Extract | Control |
| 411 | 196 | 777 | 181 | 9 | 10 |
| 544 | 39 | 808 | 615 | 8 | 11 |
| 796 | 29 | 36 | 123 | 14 | 16 |
| 553 | 39 | 499 | 595 | 12 | 10 |
| 338 | 27 | 181 | 615 | 21 | 9 |
| 156 | 796 | 544 | 540 | 21 | 7 |
| 102 | 481 | 527 | 83 | 3 | 10 |
| 798 | 449 | 492 | 333 | 2 | 11 |
| 665 | 113 | 316 | 623 | 10 | 7 |
| 410 | 609 | 384 | 261 | 13 | 15 |

**Raw Data for Figure S3**

| Ether Ex. | Control | MeOH Ex. | Control | BuOH Ex. | Control |
|---|---|---|---|---|---|
| 1 | 6 | 0 | 17 | 2 | 25 |
| 5 | 13 | 1 | 21 | 1 | 25 |
| 2 | 23 | 2 | 9 | 2 | 17 |
| 0 | 5 | 2 | 4 | 3 | 11 |
| 10 | 16 | 6 | 46 | 2 | 12 |
| 0 | 11 | 1 | 41 | 7 | 14 |
